# Unraveling Environmental Drivers of Insect Pest Dynamics Across Multiple Landscapes

**DOI:** 10.64898/2026.09.16.752066

**Authors:** Cécile Caumette, Marie-Pierre Chapuis, Sylvain Piry, Thierry Brévault, Ousmane Ndoye, Aristide Diatta, Julien Papaïx, Karine Berthier

## Abstract

The persistence of pest populations during crop off-seasons is a major challenge for integrated pest management, yet the ecological processes underlying survival and re-infestation often remain unclear. We investigated the oriental fruit fly, *Bactrocera dorsalis*, a major mango pest, through a two-year demographic monitoring across 56 orchards in the two main mango production basins in Senegal: the Sahelian Niayes and the tropical Casamance. Using a framework combining mechanistic and machine learning models, we identified the main environmental drivers of two key demographic parameters: the local abundance during the mango off-season and the onset of population growth in orchards. In the Niayes, harsh dry-season conditions confined residual populations to micro-habitats providing humidity and shade, while alternative hosts such as citrus trees could act as catalysts for early outbreaks. In contrast, under tropical conditions in the Casamance basin, *B. dorsalis* populations were more abundant year-round, and higher off-season densities appear to delay outbreak onset, likely due to density-dependent processes such as competition and predation. These findings show that distinct ecological processes shape *B. dorsalis* off-season persistence across environments, thereby determining population carry-over and growth onset, and ultimately leading to contrasting population dynamics across environments. They support region-specific management strategies to effectively reduce pest pressure: targeting refuges and early host resources in the Niayes, and enhancing natural regulation in Casamance.

## Introduction

During the crop off-season, crop pests experience drastic reductions in resource availability, as well as unfavourable environmental conditions, due to low winter temperatures in temperate regions or the dry season in arid and tropical areas. Population persistence may then rely on different processes, including local dispersal to find favourable refuges (Hickmann et al. 2025; Kennedy and Storer 2000). A given habitat may act as a refuge for pest populations by supporting several key biological functions, which are not mutually exclusive, such as feeding (e.g., presence of alternative hosts that provide resources outside of crop seasons), reproduction (e.g., availability of fruiting hosts that allow oviposition and larval development), and survival under otherwise unfavourable conditions (e.g., microclimatic shelters that buffer extreme temperatures). Identifying habitat features that allow local persistence of pest populations during the off-season is then fundamental to understand crop infestation dynamics and design effective integrated pest management. For example, specifically targeting off-season habitats sheltering residual pest populations provides an opportunity for deploying early interventions to suppress the sources of infestation before the cropping season. However, their assessment is notoriously difficult, as they can occur in non-plant environments (e.g., dormant stages in the ground), non-host plants (e.g., providing favourable microclimates), or alternative plant hosts, located within or outside crop fields and possibly scattered across the surrounding landscape (Hickmann et al. 2025; Rusch et al. 2010; Saeed et al. 2015). In this context, identifying and characterizing crop pest refuges requires landscape-scale approaches encompassing broad spatial extents, such as entire production basins, extensive sampling efforts to collect demographic data, such as abundance, and a robust statistical framework to relate demography to environmental characteristics. Furthermore, mechanistic insights into the ecological processes driving pest spatio-temporal dynamics can be gained by estimating key population parameters, such as temporal changes in population size, rather than relying on raw demographic measures. For example, in a recent study on the oriental fruit fly, *Bactrocera dorsalis* (Hendel, 1912) (Diptera: Tephritidae), Caumette et al. (2024) proposed an analytical framework combining a mechanistic demographic model to infer the annual onset of infestations in crop fields with a machine learning algorithm relating spatiotemporal variation in this parameter to environmental predictors from field to regional scales. This approach can be extended to jointly estimate additional demographic parameters across different phases of population dynamics, such as the abundance of residual populations during the off-season. Such estimates may facilitate discrimination among alternative hypotheses regarding the relative contributions of key ecological processes, such as local persistence and immigration from source populations, to annual crop reinfestation.

Furthermore, access to these refuges strongly depends on the interplay between pest life-history traits, notably dispersal abilities, and environmental heterogeneity, including landscape structure, agricultural practices, communities of natural enemies, and climatic factors. Therefore, even for a single species, population dynamics may vary considerably across landscapes, reflecting context-dependent interactions between pest populations and environmental conditions across spatial and temporal scales (Gutiérrez Illán et al. 2020). In this context, conducting landscape-scale comparative studies across contrasting production basins is essential to capture species–environment interactions, understand the mechanisms driving population dynamics, and design effective, context-specific management strategies (Brévault and Clouvel 2019). However, such approaches require intensive and consistent demographic data collection across space and time, along with the construction and harmonization of datasets on land cover, climate, and agricultural practices at comparable spatial scales, so as to ensure accuracy, robustness and interpretability of the results (Redlich et al. 2022; Warton et al. 2015). Overall, despite their promise, integrative landscape-scale approaches remain resource-intensive and methodologically demanding, and are, in practice, rarely implemented, even less so across contrasting environments.

In the present study, we address this challenge by focusing on the oriental fruit fly, *Bactrocera dorsalis*, as a model system. This invasive species, native to tropical Asia, has emerged as a major pest of mango and other tropical fruit crops in Africa in the early 2000s (Ekesi et al. 2006). Larval feeding within fruits and quarantine restrictions in European markets cause substantial economic losses to the mango industry (Ekesi et al. 2011; Mutamiswa et al. 2021; Ndiaye et al. 2024; Vayssières et al. 2008). Although *B. dorsalis* is highly polyphagous, with a broad host range including both cultivated and wild plants, mango remains its preferred host in Africa (Drew et al. 2005; Ekesi et al. 2006; Motswagole et al. 2019; Vayssières et al. 2009). We focused on the two main mango production basins in Senegal: the Niayes, a north-western coastal region near Dakar, and the Basse-Casamance region in the extreme southwest of the country. These two landscapes span a strong bioclimatic gradient, from Sahelian to tropical conditions, and exhibit markedly contrasting spatiotemporal dynamics of BD populations. In the Niayes, *B. dorsalis* abundance shows a sharp seasonal peak during the mango season followed by a drastic demographic collapse, leading to near-zero abundance during the dry season (Caumette et al. 2024; Dieng et al. 2019). In contrast, populations in Basse-Casamance persist at higher levels throughout the year, while still exhibiting a well-pronounced seasonal peak during mango fruiting, a pattern that has been associated with more humid conditions and the availability of diverse alternative host plants (Dieng et al. 2019; Konta et al. 2016). (Dieng et al. 2019; Konta et al. 2016).

We conducted standardized monitoring of *B. dorsalis* abundance over two consecutive annual population cycles, in 56 orchards, selected to capture environmental heterogeneity and ensure broad spatial coverage. We complemented these data with multi-scale environmental descriptors of cropping systems, landscape structure, and climatic variability across both regions (∼5,000 km^2^ each). Building on the work of Caumette et al. (2024), we applied a similar analytical framework combining demographic inference and machine learning to identify the main environmental drivers of population dynamics. In this framework, we adapted a mechanistic demographic model (Soulsby and Thomas 2012) to jointly estimate two parameters associated with different seasonal phases of the population cycle: (i) off-season residual abundance and (ii) the onset of population growth in orchards. This dual-landscape approach, which provides a natural contrast between two environments influencing *B. dorsalis* population dynamics, allows us to test the following hypotheses. Under the harsh environmental constraints of the Sahelian Niayes basin, both demographic parameters, residual population abundance and the onset of population growth, are expected to be primarily driven by local habitat conditions that promote persistence, for example by buffering thermal and hydric stress and providing access to scarce early fruit resources when available. In particular, refuge habitats with larger residual populations are expected to promote earlier population growth and act as sources for the re-infestation of orchards at the onset of the mango season (Caumette et al. 2024). Such knowledge could inform the temporal and spatial targeting of early-season interventions aimed at reducing pest pressure in refuge habitats before the onset of the mango season. In contrast, in the tropical, moist Basse-Casamance basin, mild climatic conditions and high host plant diversity are expected to promote larger residual populations through higher survival and continuous reproduction throughout the mango off-season, leading to reduced dependence on specific local habitats and earlier population growth (Dieng et al. 2019; Konta et al. 2016). The fine-tuned, targeted management approach proposed for the Niayes is unlikely to be transferable to such contexts, where management should instead prioritize broader strategies aimed at reducing overall reproduction and survival, for example by promoting natural enemies.

## Material and Methods

### Study areas and monitoring of *B. dorsalis* abundance data

The two study areas, of about 5000 km^2^ each, are located in the two main Senegalese mango production basins: the Niayes, located northwestern Senegal, near the city of Dakar, and the Basse-Casamance, located in the southwestern region of Senegal near the border with Guinea-Bissau (Fig. 1A). The Niayes area is under a Sahelian climate with a short rainy season from July to September (around 400-600 mm rainfall) followed by a long dry season from October to June (Diallo et al. 2015; Grechi et al. 2013; Vayssières et al. 2011). This is the major area of fruit and vegetable production in Senegal and mango is the main fruit produced for international and national markets as well as local consumption (Dieng et al. 2019; Grechi et al. 2013; Vayssières et al. 2011). Landscape otherwise alternates between cultivated lands, urban areas and scarce natural vegetation (Fig. 1A). In contrast, the Basse-Casamance region (hereafter referred as to Casamance) is closer to the species’ native range, with a tropical Sub-Guinean climate with a heavy rainy season from May to October with more than 1200 mm of rainfall (Dieng et al. 2019; Traore et al. 2017). Crossed by the Casamance River, the landscape is a mosaic of dense forest, mangrove and paddy fields (Andrieu and Mering 2008). Mangoes are mainly produced in familiar and traditional orchards scattered across the landscape and are dedicated to local or national consumption (Ba et al. 2020). The Casamance area is also the cashew nut basin of Senegal and the country has doubled the surface production of this fruit between 1986 and 2017 (Cabral and Costa 2017; Ndiaye et al. 2020). Infestations of cashew apples, the fleshy false fruit of the cashew tree, have been reported in Africa and Asia, and although it is considered largely secondary to mango trees, cashew is listed as a host plant of BD in the European and Mediterranean Plant Protection Organization (EPPO) Global Database (https://gd.eppo.int/).

**Figure 1:**
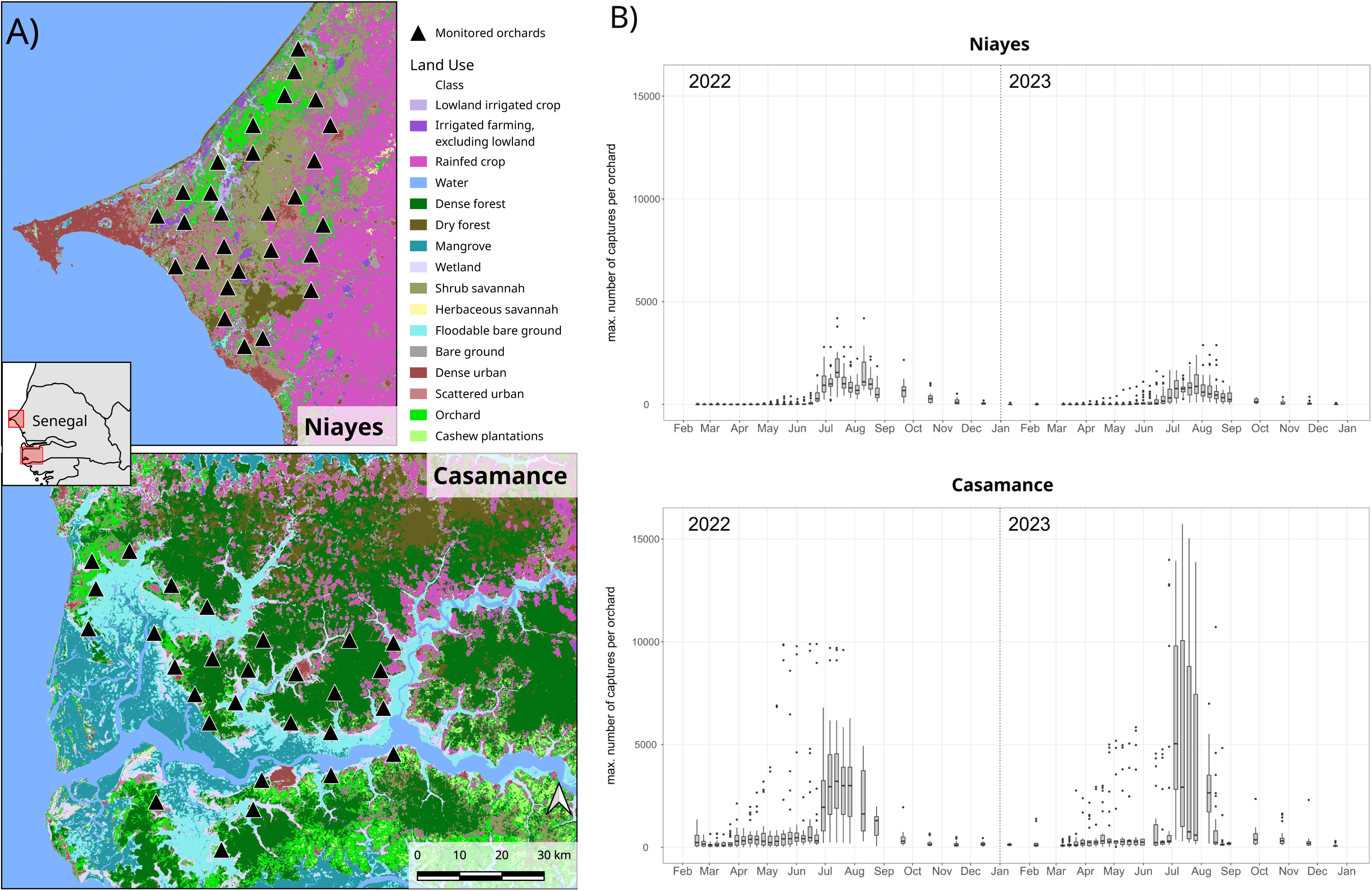
Localisation of monitored orchards (A) and temporal variation of *B. dorsalis* abundance (B) in the two main mango production basins in Senegal, the Niayes (upper panel) and Casamance (lower panel). (A) The background map represents the 16 land cover classes considered in this study (see Supplementary material for detail). (B) Abundance data was recorded weekly (March to August) or monthly (September to February) over two years of monitoring. Each boxplot shows the median value (black line), the lower (Q1) and upper (Q3) quartiles (lower and upper box limits), the highest and lowest values excluding outliers (vertical lines, with a maximum length of 1.5*(Q3-Q1)) and outliers (black dots) of the maximum number of flies trapped.

The monitoring of *B. dorsalis* abundance was conducted from February 2022 to august 2024 within 28 orchards within each production basin. Orchards were selected in order to maximize both the spatial coverage and the representation of the environmental diversity (Fig. 1A). The monitoring protocol was similar to the one described in Caumette et al. (2024). In brief, three traps were set up in each orchard and baited with methyl-eugenol, an attractant for *B. dorsalis* males recommended for early detection and estimation of *B. dorsalis* abundance (Manrakhan et al. 2017; Manrakhan et al. 2019). Traps were collected weekly from March to August and monthly from September to February, with a one-week exposure period in all cases to ensure comparable sampling effort throughout the year. For each orchard and sampling date, the number of flies caught in each trap was counted.

### Estimation of *B. dorsalis* population demographic parameters

Using the abundance data collected in 2022 and 2023 in the Niayes and Casamance areas (Fig. 1B), we built annual abundance times series for each orchard by considering the maximum number of flies caught among the three traps at each sampling date (see Supplementary Material S1). We then applied a modified version of the POPFIT mechanistic model (Soulsby and Thomas 2012). The model was adapted by relaxing the assumption that baseline abundance (i.e. before the onset of population growth and after the seasonal decline at the end of the year) is zero, allowing for non-zero residual populations in the fitted growth curves (see Supplementary Material S2). This framework allowed us to estimate two demographic parameters for each orchard and year: (i) the onset of population growth (*t*_0_), as in Caumette et al. (2024), and (ii) mean abundance during the low-density period (*eps*). Parameter inference was conducted using a mechanistic-statistical approach (Papaïx et al. 2022), within a Bayesian framework, implemented in the R package *nimble* v. 1.2.1 (de Valpine et al. 2017) (see Supplementary Material S2, for details on the parameters and their prior distributions). For each demographic parameter, we obtained a posterior distribution for every orchard-year combination (112 in total), representing the range of plausible values given the data. We randomly sampled 500 values from each distribution to build 500 distinct sample sets, each containing one randomly selected value per orchard–year combination. This approach allowed us to account for uncertainty in parameter estimation in subsequent analyses assessing the influence of environmental variables (see Supplementary Material S5 in Caumette et al. (2024) for details on the construction of the sample sets).

### Environmental predictors and modeling of their effects on population parameters

An overview of the multiscale environmental predictors describing the cropping system, the landscape structure and the weather variability is provided in Table 1, and details on their construction and preprocessing are given in Supplementary Material S3. Beyond processing heterogeneous data sources for adaptation to our country-scale production-basin framework, we improved the characterization of environmental predictors in three main ways Caumette et al. (2024) : (i) the development of a refined 16-class land cover typology tailored to study *B. dorsalis* population ecology and incorporating classification uncertainty; (ii) the use of the R package *scalescape* v. 0.0.0.9 (Lowe et al. 2022) to estimate spatial ranges of landscape influence on demographic parameters, extending the *siland* framework (Carpentier and Martin 2021; Caumette et al. 2024) by enabling the joint estimation of spatial influence functions across all classes; and (iii) a refined characterization of orchard host resources based on phenology-defined classes, explicitly accounting for differences in fruiting periods and their relative frequencies.

**Table 1:**
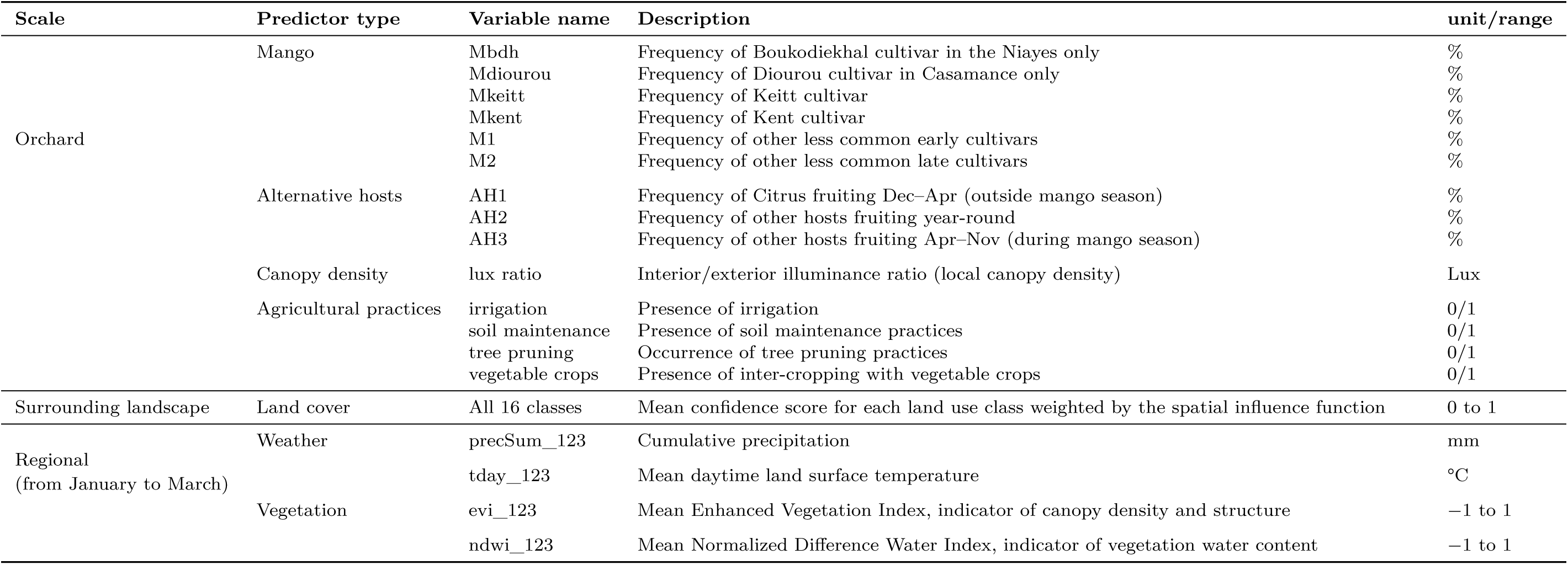
The 34 multi-scale environmental predictors included in the models used to assess effects on population parameters. The land cover nomenclature is provided in Supplementary Material S3.

| Scale | Predictor type | Variable name | Description | unit/range |
| --- | --- | --- | --- | --- |
| Orchard | Mango | Mbdh | Frequency of Boukodiekhall cultivar in the Niayes only | % |
|  |  | Mdiourou | Frequency of Diourou cultivar in Casamance only | % |
|  |  | Mkeitt | Frequency of Keitt cultivar | % |
|  |  | Mkent | Frequency of Kent cultivar | % |
|  |  | M1 | Frequency of other less common early cultivars | % |
|  |  | M2 | Frequency of other less common late cultivars | % |
|  | Alternative hosts | AH1 | Frequency of Citrus fruiting Dec–Apr (outside mango season) | % |
|  |  | AH2 | Frequency of other hosts fruiting year-round | % |
|  |  | AH3 | Frequency of other hosts fruiting Apr–Nov (during mango season) | % |
|  | Canopy density | lux ratio | Interior/exterior illuminance ratio (local canopy density) | Lux |
|  | Agricultural practices | irrigation | Presence of irrigation | 0/1 |
|  |  | soil maintenance | Presence of soil maintenance practices | 0/1 |
|  |  | tree pruning | Occurrence of tree pruning practices | 0/1 |
|  |  | vegetable crops | Presence of inter-cropping with vegetable crops | 0/1 |
| Surrounding landscape | Land cover | All 16 classes | Mean confidence score for each land use class weighted by the spatial influence function | 0 to 1 |
| Regional<br>(from January to March) | Weather | precSum_123 | Cumulative precipitation | mm |
|  |  | tday_123 | Mean daytime land surface temperature | °C |
|  | Vegetation | evi_123 | Mean Enhanced Vegetation Index, indicator of canopy density and structure | –1 to 1 |
|  |  | ndwi_123 | Mean Normalized Difference Water Index, indicator of vegetation water content | –1 to 1 |

Following Caumette et al. (2024), we used the tree-boosting method GPBoost (Sigrist 2022), implemented in the R package *gpboost* v. 1.4.0.1 (Sigrist et al. 2023), to hierarchize the effects of 34 multi-scale environmental predictors on local abundance during the mango off-season (*eps*) and the onset of demographic growth (*t*_0_), within each of the two study regions. A full description of the GPBoost analytical framework is provided in Supplementary Material S5 of Caumette et al. (2024), and details of model testing are provided in Supplementary Material S4. Briefly, all environmental candidate factors were considered as fixed effects and the year of sampling as a grouped random effect in the final GPBoost model. In addition, for the *t*_0_ parameter, *eps* was also considered as a predictor in the model. For each demographic parameter and each production basin, we performed 500 independent analyses on the 500 demographic sample sets. The overall contribution of a given predictor to the model output (hereafter called *S_mean_*) was obtained by averaging the absolute SHAP values of the observations. For each demographic parameter and basin, the predictors were ranked in decreasing order based on the median of the *S_mean_* values across the 500 analyses. Based on this ranking, the relationship of each of the most important predictors with the estimated parameters *eps* and *t*_0_ within each study area was investigated using a dependence plot built by fitting individual SHAP values from the 500 analyses as a gam-smoothed function of the predictor values, using the R package *mgcv* v. 1.9-1 (Wood 2017). Finally, as a validation step of the variable selection procedure, we assessed the performance of the GPBoost models in predicting *eps* and *t*_0_ in each study area by randomly partitioning each sample set into a training dataset and a test dataset, i.e. 80% and 20% of the data, respectively. Model accuracy was then assessed by computing the Pearson correlation coefficient between predicted and observed values as well as the Root Mean Square Error (RMSE) of the model for each sample set.

## Results

The Bayesian estimation of population residual abundance and onset of growth, *eps* and *t*_0_, correctly converged for all annual time series, with Gelman–Rubin statistic values *≤* 1.1. Parameter estimates showed high precision, with standard deviations of posterior distributions *≤* 0.26 week for *t*_0_ in both production basins, and *≤* 3.2 and 10 individuals for *eps* in the Niayes and Casamance production basins, respectively (Table 2). Consistent with previous studies (Caumette et al. 2024; Dieng et al. 2019), abundance time series showed clear differences in *B. dorsalis* population dynamics between the Niayes and Casamance basins (Figures 1B and 2), with highly significant Wilcoxon rank-sum tests between basins for both population parameters and both years. In the Niayes, *B. dorsalis* populations were nearly absent outside the demographic peak, with mean abundance across orchards reaching only a few individuals in both years. In contrast, in Casamance, mean abundance during the low demographic phase reached hundreds of individuals, indicating that *B. dorsalis* remained relatively abundant during the off-season in 2022 and 2023. In both basins, the mean onset date of within-orchard population outbreaks (in weeks from 1^st^ January) occurred in May (Wilcoxon rank-sum test *p*-value between basins: 0.392 in 2022 and 0.001 in 2023) (Table 2). However, in Casamance, more orchards were infested early (January–February), and the interval between the earliest and latest *t*_0_ values was longer (22 and 28 weeks in 2022 and 2023, respectively) than in the Niayes (13 and 17 weeks in 2022 and 2023, respectively) (Table 2).

**Figure 2:**
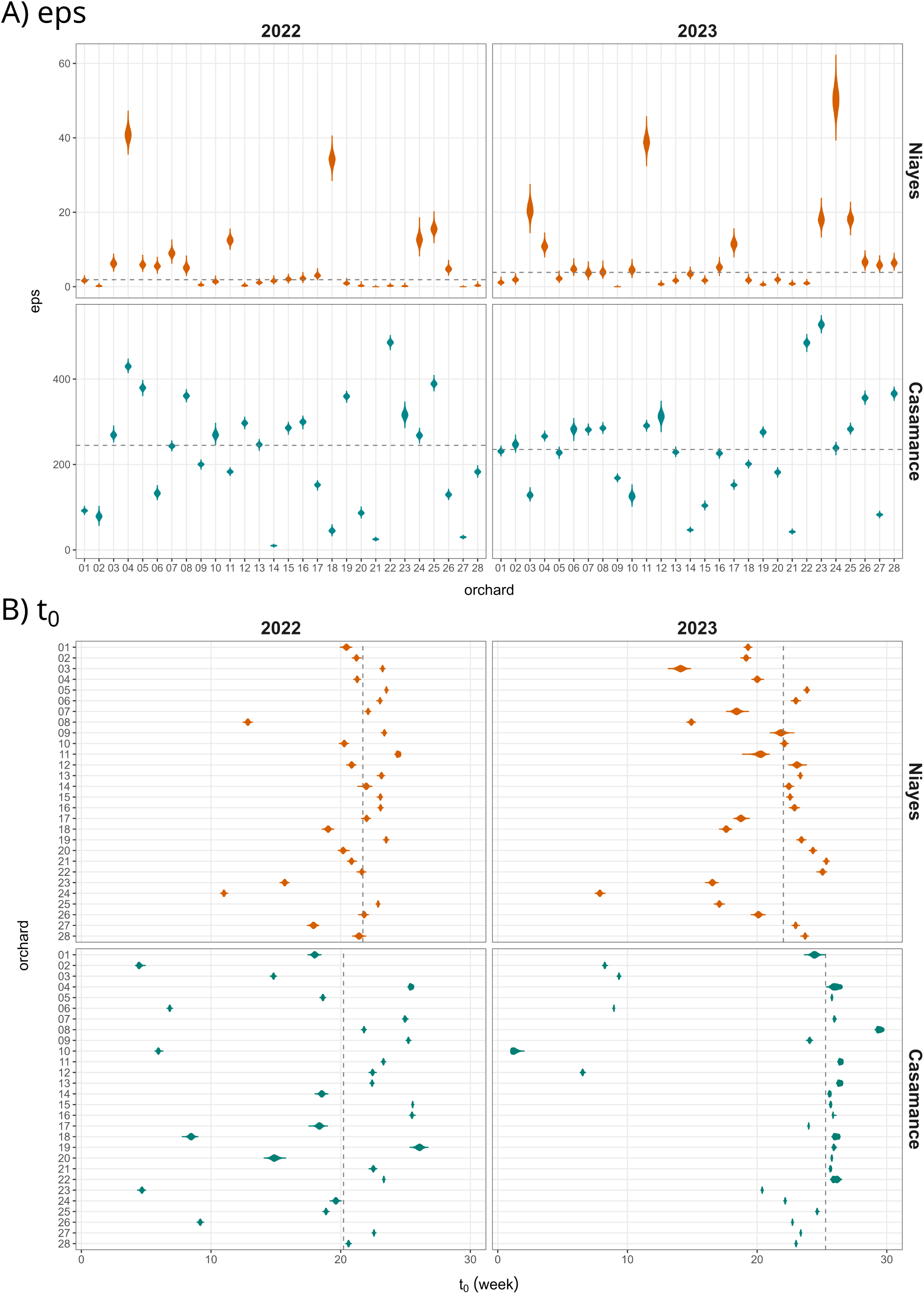
POPFIT Bayesian posterior distributions of the demographic parameters per orchard and year within both production basins. For each combination of year and orchard in the Niayes (top panel) or in Casamance (bottom panel), the violin plot represents the posterior distribution of A) *eps*, the *B. dorsalis* abundance level during the off-season (NB: here, for the sake of clarity, the scale is different for the two basins), B) *t*_0_, the onset of *B. dorsalis* population growth. Horizontal, respectively vertical, dashed lines represent the median values of the parameters.

**Table 2:**
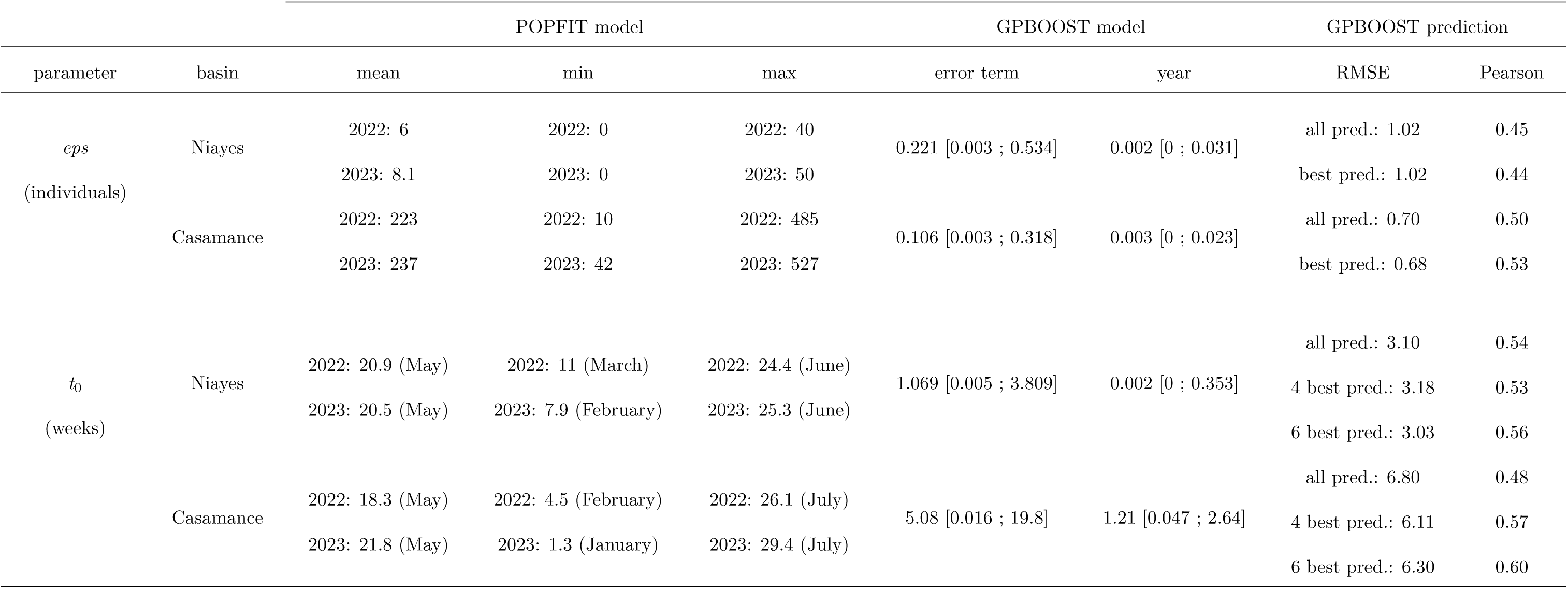
Overview of results from POPFIT and GPBoost models. . POPFIT: summary statistics (mean, minimum, and maximum) of the Bayesian posterior distributions across the 28 orchards are reported for each demographic parameter, production basin, and year. GPBoost: for each parameter and production basin, the table shows the model’s error term, year random effect, and validation metrics from the prediction (mean of RMSE and Pearson correlation over the 500 replicates) calculated using either all predictors or the top-ranked ones (indicated as "all" or "best" in the table)

For each demographic parameter and production basin, the SHAP-based ranking of the environmental predictors from the GPBoost models is presented in Figure 3. The effects of the top ranked predictors on the demographic parameters within each basin are presented in Figure 4. In the Niayes basin, a group of six predictors clearly emerge as important in explaining *B. dorsalis* abundance during the low demographic phase (*eps*), with the presence of vegetable crops within orchards being by far the strongest predictor, with a positive effect and an average SHAP value of around 0.28. The mean of the SHAP values of the next five predictors was very similar, around 0.1. Among them, the positive effect of *NDWI* and *precSum_123* on *eps* suggested that higher levels of humidity favoured the maintenance of local *B. dorsalis* populations during the dry period. The remaining three predictors were local cropping system variables: denser tree cover (lux ratio *<* 0.25), a high proportion of the mango cultivar Boukodiekhal (*>* 50%), and a low proportion of the Kent cultivar (*<* 25%) were associated with higher abundance of *B. dorsalis* during the mango off-season. In the Casamance basin, the ranking and the relationships of the environmental predictors with *eps* was less clear, with three of the top predictors suggesting local or confounding effects (i.e., two orchards driving the negative relationships observed for the landscape classes *shrub savannah* and *floodable bare ground* and negative impact associated to the strict absence of the landscape class *rainfed crop*). The three other top predictors included the proportions of mango and cashew orchards in the surrounding landscape, with higher proportions of both associated with lower fly abundance during the off-season. The level of precipitation (*precSum_123*) was also identified as an important predictor and, similarly to the Niayes basin, showed a slight positive effect on *B. dorsalis* abundance during the low demographic phase.

**Figure 3:**
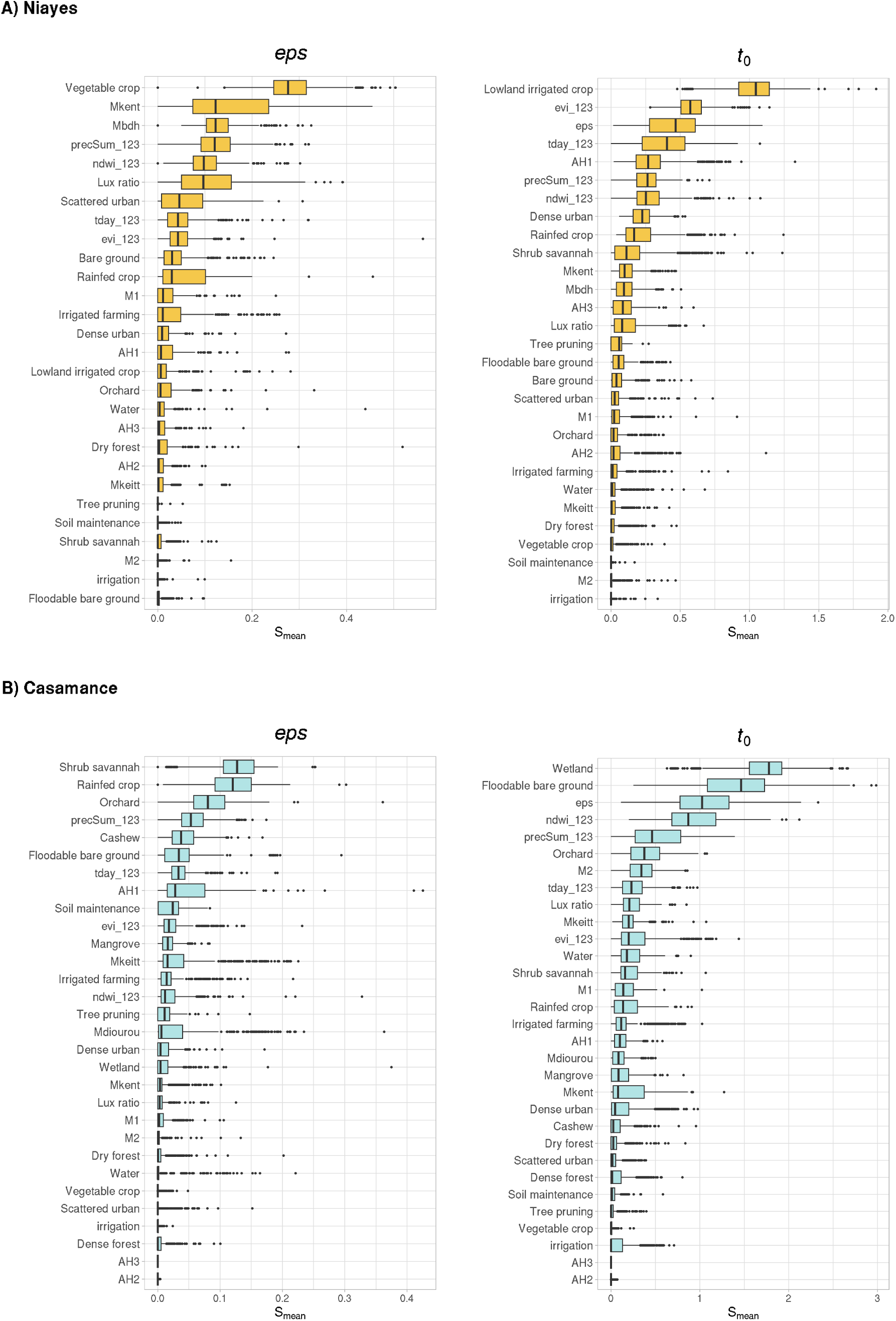
Ranking of the environmental predictors of *B. dorsalis* abundance during the mango off-season (*eps*; left panel) and onset of population growth (*t*_0_; right panel) in the A) Niayes and. . **B) Casamance basins.** Ranking is based on the SHAP values of the predictors resulting from the GPBoost model applied independently to the 500 sample sets for each parameter and production basin. Boxplots show median, first and third quartiles, range excluding outliers (whiskers), and outliers (black dots).

**Figure 4:**
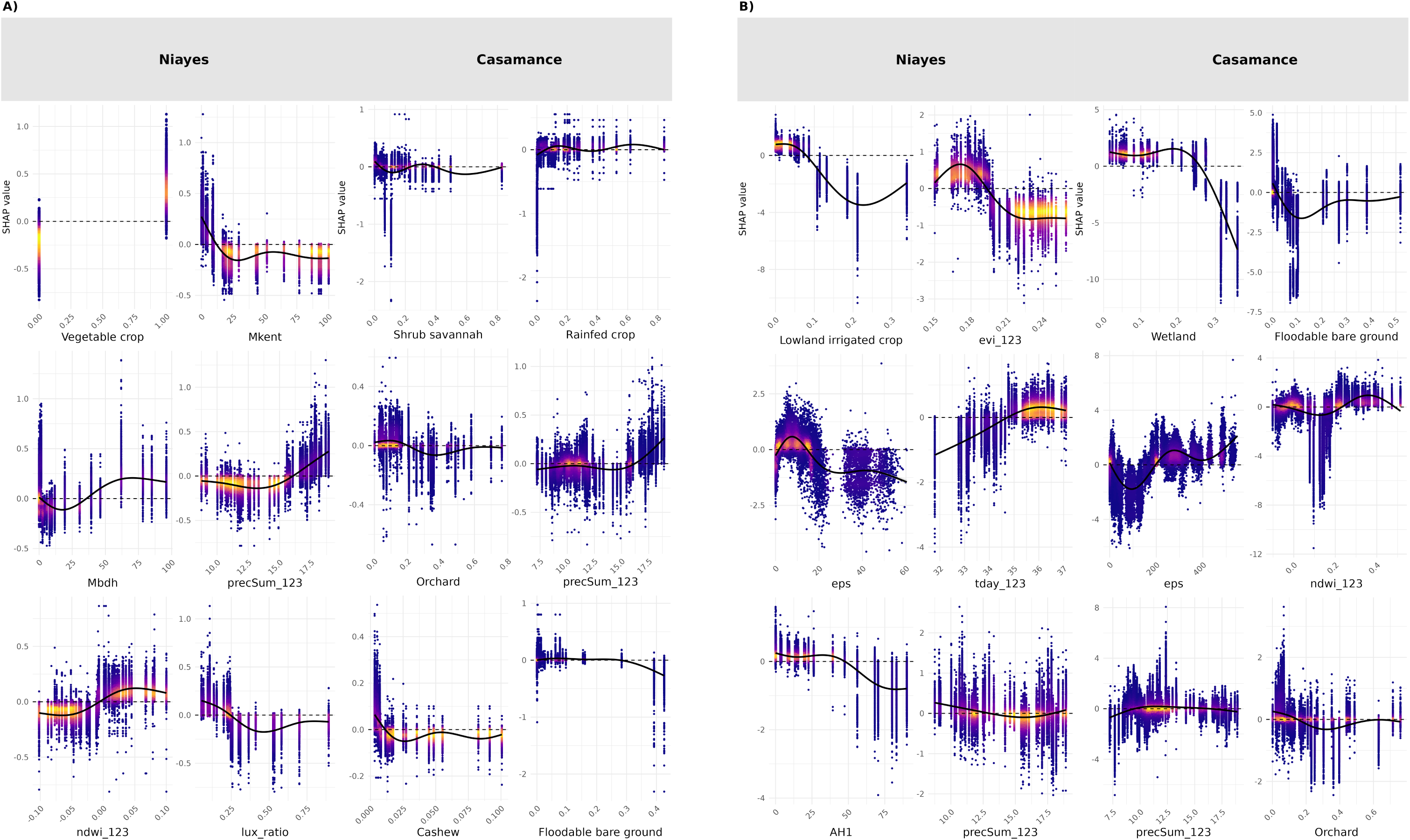
SHAP dependence plots of the top environmental predictors selected from the GPBoost model for A) *eps* and B) *t*_0_, in the Niayes and Casamance basins. For quantitative predictors (all except *vegetable_crop* in the Niayes), the GAM-smoothed curve of SHAP values across the 500 sample sets is shown as a black line and the density of SHAP values is displayed with a gradient color from high density in yellow to low density in violet. The x-axis corresponds to the predictor values. GAM were fitted using thin plate regression splines and by fixing the basis dimension *k* to 5 to avoid overfitting. For *vegetable crop* (qualitative predictor), the graph shows the distribution of the SHAP values over the 500 sample sets according to the presence (x-axis value = 1) and absence of vegetable crops (x-axis value = 0) in the orchards.

For each production basin, a group of four predictors stands out as the most meaningful to explain the variability in the annual onset of *B. dorsalis* populations within orchards (*t*_0_). In both basins, the abundance of local populations (*eps*) during the mango off-season is a clear predictor of the timing of population growth in orchards, although the effect is opposite in each basin. In the Niayes, the presence of at least a few dozen of individuals were associated to early outbreak within orchards. In contrast, in Casamance, the local presence of large *B. dorsalis* populations during the mango off-season (e.g., several hundred individuals) clearly delayed the onset of the outbreak phase in orchards. In the Niayes, very early population growth was strongly associated with lowlands, which are wet depressions where groundwater is generally close to the surface and that are used as small, intensive, and often highly diversified horticultural plots (Sall and Vanclooster (2009)). The favourable conditions that can be found in these environments are supported by other top-ranked predictors: early outbreak was promoted when *B. dorsalis* was sheltered from temperatures exceeding 35°C by a dense canopy, as indicated by the contrasting effects of *EVI* (negative effect on *t*_0_) and *tday_123* (positive effect on *t*_0_), and when citrus trees constituted at least 50% of the orchard and the level of occasional rainfall during the dry season (*precSum_123*) was higher (i.e., negative effect of both predictors on on *t*_0_). Interestingly, in the Casamance basin, the ranking of the environmental predictors of *t*_0_ indicates that the presence of certain seasonally wet habitats may favour early population growth, as suggested by the negative relationship between *t*_0_ and the landscape classes *wetland* and *floodable bare ground*. However, excessive humidity, associated with vegetation or precipitation, delayed orchard infestation, as indicated by the positive relationship between *t*_0_ and the variables *NDWI* and *precSum_123*.

Finally, the predictive performance of reduced GPBoost models based on the top-ranked predictors was comparable to that of the full model including all 34 candidate predictors. The reduced model included the six top-ranked predictors for *eps* and either the four or six top-ranked predictors for *t*_0_. The median values and distributions of the RMSE and Pearson correlation coefficient across the 500 independent analyses of the sample sets were very similar (see Table 2 and Supplementary Material S5), siggesting that the reduced set of top-ranked predictors captures much of the relevant environmental signal, while reducing the number of less informative predictors and potentially limiting noise.

## Discussion

In this work, we applied the analytical framework proposed by Caumette et al. (2024) to hierarchize the effects of multi-scale candidate environmental factors on the seasonal population dynamics of *B. dorsalis*, a major pest of mango crops in Senegal. By combining mechanistic demographic inference and machine learning methods, this framework integrates process-based and correlative approaches within a single analytical workflow, explicitly accounting for spatio-temporal structure in population processes. In addition, the joint estimation of residual population abundance and onset of population growth enables the exploration of different phases of the seasonal dynamics within an unified framework. This dual perspective is particularly suited to disentangling context-dependent ecological processes across heterogeneous landscapes. Applied to contrasted environmental contexts such as the Niayes and Casamance production basins, this comparative design provides insights into distinct density-dependent ecological processes in the two regions. Our results, detailed below, should nevertheless be interpreted with caution, acknowledging several limitations. First, despite extensive spatial coverage, some environmental and landscape categories remain unevenly represented, particularly in the Casamance basin, which may reduce the ability to detect weak or rare effects. In addition, while the modelling framework captures complex non-linear relationships and interactions, it remains correlative and therefore does not allow formal inference of causality. Consequently, our results are also discussed in light of ecological processes that are not explicitly represented in the models but may be indirectly reflected by the selected environmental predictors.

### Contrasting population dynamics across production basins

An interesting result is the apparent opposite relationship between the abundance of *B. dorsalis* during the mango off-season and the timing of the demographic growth within orchards across the Niayes and Casamance basins. This pattern may reflect a consistent density-dependent relationship expressed across different portions of the population density gradient between the two basins. In the Niayes, residual populations of just a few dozen individuals are associated with the earliest onsets of population growth. In contrast, in Casamance, where population abundance during the low demographic phase is generally higher by a factor of 10, the demographic peak tends to be delayed when local populations already reach several hundred individuals during the off-season. This difference in seasonal dynamics may result from both direct and delayed density-dependent effects. High population density in a given habitat during the less favourable season can immediately increase mortality through predator aggregation (Eveleigh et al. 2007), and reduce reproductive success due to competition for limited oviposition sites, as fruit availability is still low and females of *B. dorsalis* typically avoid laying eggs in fruits already infested by conspecific larvae (Clarke 2019; Fletcher 1987). In addition, high densities may limit immigration from surrounding populations due to overcrowding and unfavourable local conditions (Clobert et al. 2009). Resource limitation and physiological stress can also reduce adult condition or fecundity in the following generation (Reyes-Ramírez et al. 2023). Such processes could delay population outbreaks in Casamance orchards, where *B. dorsalis* densities remain high during the mango off-season, causing the main population increase to occur only when resources become highly abundant, i.e., at the peak of the mango season. In contrast, in the Niayes, local residual populations are very small when present, likely resulting in reduced predation risk due to limited predator attraction and weaker resource competition early in the mango season, followed by subsequent immigration as mangoes become increasingly available for oviposition.

### Determinants of off-season abundance and persistence

Differences in *B. dorsalis* abundance during the mango off-season between the two basins have already been linked to the contrasted bioclimatic conditions, with the Casamance being a highly forested area, with a potentially large range of alternative hosts and a tropical climate similar to the native range of *B. dorsalis*, while the Niayes is basically an unfavourable environment characterized by a Sudano-Sahelian climate and sparse natural vegetation (Konta et al. 2016). Consistently, and in line with the results of Caumette et al. (2024), we found that in the Niayes, the population abundance during the off-season was positively associated to highest levels of humidity, associated to vegetation (*NDWI_123*) or precipitation events (*precSum_123*), and local tree shading (*lux ratio*). These results support the hypothesis of Caumette et al. (2024) that, in the Niayes, favourable microclimate conditions provide refuges for small populations that could survive, with little or no reproduction during the mango off-season. Interestingly, in the present study, which is based on a better representation of the environmental heterogeneity in the Niayes (from the orchard to the basin scale), we found that the main predictor of local *B. dorsalis* abundance during the off-season was the presence of vegetable crops within orchards. Although fruits (e.g. mango, citrus, guava) are identified as the preferred hosts for *B. dorsalis*, vegetable crops such as tomatoes, peppers, and cucurbits have also been reported as potential hosts for larval development (African host plants reviewed in Mutamiswa et al. (2021)). Reports of larval emergence from vegetable crops in the Niayes (Boinahadji et al. 2019; Ndiaye et al. 2012) suggest that their presence in some orchards may provide a small but potentially sufficient resource supply to support limited reproduction. However, a monitoring survey of Solanaceae and Cucurbitaceae crops conducted in 2012–2013 in the Niayes did not detect any infestation by *B. dorsalis* (Brévault, pers. comm.), suggesting that the presence of vegetable crops may instead reflect highly favorable microclimatic conditions that support *B. dorsalis* persistence throughout the dry season, as they are regularly watered from local water sources, such as wells or wet depressions.

The results of the GPBoost model also indicate additional effects of the cropping system, specifically the proportion of mango cultivars Kent and Boucodiekhal (BDH), with a negative effect and positive effect, respectively, on population abundance during the off-season. These opposite effects may indirectly result from differences in orchard management and structure, which likely shape microclimatic conditions and habitat suitability for the persistence of residual populations during this period. In the Niayes, the Kent cultivar dominates intensive, mono-specific orchards dedicated to the export market. These orchards are strictly managed, including practices such as tree pruning, pesticide application, soil maintenance, removal of fallen and aborted mangoes, and early harvesting for export (Grechi et al. 2013; Ndiaye et al. 2024; Ndiaye et al. 2012). Such intensive management is likely to greatly reduce the presence of *B. dorsalis* during the dry season. In contrast, BDH is primarily grown in traditional, low-input orchards intended for local consumption, where management is minimal, with typically no regular pruning, no removal of aborted or fallen fruits, and a more extended harvest period depending on household needs (Grechi et al. 2013; Ndiaye et al. 2012). The resulting high tree cover, along with the availability of fallen, aborted, and unharvested fruits, likely creates favourable micro-habitats that support *B. dorsalis* persistence.

In the Casamance basin, among the six top predictors of *B. dorsalis* abundance during the off-season, only the level of precipitation showed a positive effect, although less pronounced than in the Niayes. Other key predictors included the proportion of mango and cashew orchards in the surrounding landscape, both of which were negatively associated with local population abundance during the low phase. This pattern could reflect a dilution effect (Veres et al. 2013), where isolated orchards tend to concentrate individuals, whereas in areas with high orchard density, individuals are more dispersed. While mango orchards likely reflect the availability of primary hosts, the effect of cashew orchards, which are considered minor hosts, may instead capture broader landscape structure effects, as increasing cashew dominance can coincide with reduced availability of mango, in some areas of Casamance. The interpretation of the other three top predictors of *B. dorsalis* abundance during the off-season is less straightforward, as they likely capture indirect or confounded effects driven by local landscape context rather than direct ecological drivers. In particular, the relationships of shrub savannah and floodable bare ground with residual population size are driven by only a few observations. Similarly, the negative effect associated with rainfed crops appears to stem primarily from the absence of this landscape class. This threshold effect at zero may suggest that some important drivers of population abundance during the off-season are missing from our predictor set. Alternatively, orchards lacking rainfed crops in their surroundings are mostly located near the extensive mangroves along the Casamance River, which may represent less suitable habitats for *B. dorsalis*, due to the absence of suitable hosts and environmental constraints such as high salinity.

### Determinants of the onset of population outbreaks

The results on the environmental drivers of the variation in the onset of population growth (*t*_0_) in the Niayes are largely consistent with previous findings by Caumette et al. (2024). We confirmed that temperatures exceeding 35 °C, which are known to negatively affect development, survival, and fecundity traits of *B. dorsalis* (see Caumette et al. 2024), significantly delay the onset of the outbreak phase in orchards. Conversely, a high level of shading (as indicated by the EVI predictor), which favours population persistence during the dry season, promotes earlier outbreaks (see Figure 4A). An additional and noteworthy effect, not reported by Caumette et al. (2024), is the association between early population growth and a high proportion of citrus trees within orchards (above 50%). This finding suggests that citrus hosts may act as catalysts for population growth when residual populations are present but mangoes are not yet ripe for oviposition, rather than serving as true alternative hosts that sustain continuous reproduction during the off-season. Previous studies have shown that citrus fruits are primarily attractive to *B. dorsalis* for oviposition when their peel is damaged (Diatta et al. 2013; Theron et al. 2017; Theron et al. 2023). This is particularly likely toward the end of the citrus fruiting period in Senegal, in April, which coincides with the early onset of outbreaks before mangoes ripen. These conditions of higher humidity, shading and the presence of citrus are commonly found in cultivated lowlands (Sall and Vanclooster (2009)), which were clearly ranked as the main predictor of early outbreaks in the Niayes in our analysis.

In Casamance, the onset of population growth within orchards appears to be strongly influenced by humidity-related factors. The presence in the surrounding landscape of highly humid habitats tend to favour early outbreaks in orchards, particularly when the proportion of wetlands, mainly rice fields in Casamance, exceeds 30%, and the proportion of floodable bare ground is around 10%. Conversely, excessive humidity associated with vegetation, large water bodies (estimated with the predictor NDWI), and precipitation (*prec_sum123*) tends to delay the start of outbreaks in orchards. High NDWI values correspond primarily to the Casamance River and its associated mangrove ecosystems. These mangroves consist of salt-tolerant species that are unsuitable for oviposition and larval development of *B. dorsalis*. Altogether, these results suggest that outbreaks in orchards adjacent to extensive mangrove areas may be influenced more by immigration from surrounding favourable habitats than by a rapid build-up of local populations.

### Implications for landscape-level pest management

Using spatio-temporal monitoring of *B. dorsalis* abundance that optimized both spatial coverage and representation of environmental heterogeneity across the two main mango production basins in Senegal, we confirmed the findings of Caumette et al. (2024). Under the harsh environmental conditions of the Niayes, the persistence of small populations during the mango off-season is highly dependent on the presence of favourable micro-habitats providing humidity and shelter from high temperatures. The apparent importance of vegetable crops in our analysis is likely indirect, reflecting their role as a proxy for locally higher humidity levels resulting from daily watering during the dry season. Residual populations may then act as sources in the re-infestation of orchards, with citrus also likely contributing to early outbreaks when residual populations are present but mango fruits are not yet sufficiently ripe for oviposition. Altogether, these findings support the notion that early interventions in orchards, combining tree pruning and the removal of damaged and fallen mangoes and citrus, as well as the control of residual populations in habitats buffering thermal and hydric stress, could effectively reduce pest pressure prior to the onset of the mango season. Cultivated lowlands, where these favourable conditions converge, should therefore be especially targeted for early control, as they may act as foci from which outbreaks spread across the wider production basin. In Casamance, such fine-tuned, targeted management strategies are unlikely to be effective because, although important outbreaks still occur in mango orchards during the crop season, *B. dorsalis* populations exhibit more stable metapopulation dynamics, with larger residual abundances and no apparent dependence on refuge habitats in unfavourable periods of the year. Mango orchards can then act as temporary, high-quality resource patches embedded within a highly diverse landscape matrix, which may trigger density-dependent processes such as competition, increased mortality, and dispersal (Hanksi and Gaggiotti 2004; Strevens and Bonsall 2011). Limiting *B. dorsalis* populations in Casamance may therefore rely more on broader strategies aimed at reducing overall reproduction and survival during the low-density phase, such as conserving and promoting natural enemies within the landscape (Perfecto and Vandermeer 2008; Tscharntke et al. 2005). Because the tropical climate and vegetation of Casamance closely resemble those of the native range of *B. dorsalis*, such strategies may be inspired by long-established management practices in native contexts.

## Acknowledgements

This work was supported by the DISLAND and MARA projects, publicly funded by the French National Research Agency (ANR - 20-CE32-0012) and TSARA international initiative (Transforming Food and Agricultural Systems through Research in Partnership with Africa). MPC was supported by the French Agricultural Research Centre for International Development (CIRAD). CC was supported by a doctoral fellowship funded by CIRAD and ANR. KB gratefully acknowledges the support of the INRAE Department of Plant Health and Environment. Computations were performed on the INRAE BioSP computing platform, to which we are grateful for providing computational resources.

## Authors’ Contributions

MPC, SP, TB and KB designed the spatio-temporal monitoring. ON and AD conducted the weekly trap surveys. CC, SP and KB curated the data. SP and KB processed remote sensing data and produced the landscape typology. JP developed the mechanistic model, with contributions from CC, MPC and KB, and CC performed the inference of the demographic parameters. CC, SP and KB performed the statistical analyses. CC, MPC and KB wrote the first draft of the manuscript, and SP, TB and JP contributed to the final version.

## Supplementary Material

**Supplementary Material S1** – Seasonal variation in attractant efficiency

**Supplementary Material S2** – Bayesian estimation of the *eps* and *t*_0_ parameters from *B. dorsalis* abundance time series, using a modified version of the POPFIT model

**Supplementary Material S3** – Construction and preprocessing of environmental predictors

**Supplementary Material S4** – Details of model testing in the GPBoost framework

**Supplementary Material S5** – Visualisation of results on GPBoost predictive performance

## Notes

### Competing Interest Statement

The authors have declared no competing interest.

