## Supplementary Material for "Unraveling Environmental Drivers of Insect Pest Dynamics Across Multiple Landscapes"

#### Contents

**Supplementary Material S1** – Supplementary Material S1 – Seasonal variation in attractant efficiency

**Supplementary Material S2** – Bayesian estimation of the *eps* and $t_0$ parameters from *B. dorsalis* abundance time series, using a modified version of the POPFIT model

**Supplementary Material S3** – Construction and preprocessing of environmental predictors

**Supplementary Material S4** – Model testing in the GPBoost framework

**Supplementary Material S5** – GPBoost predictive performance

### Supplementary Material S1 – Seasonal variation in attractant efficiency

Population monitoring was carried out using traps baited with two commercially available methyl-eugenol attractants from different manufacturers: one supplied by Biosystemes France and the other by Econex. The two brands were systematically used in each monitored orchard. A decrease in efficiency was observed for the Econex attractant during the rainy season, see Figure S1 for an example in the Niayes basin in 2023 . The relative efficiency of the two methyl eugenol attractants was further tested using a linear mixed-effects model fitted to log-transformed weekly trap catches collected over the two years. The model included attractant type, season, and their interaction as fixed effects, and orchard ID as a random effect. The rainy season was defined according to basin-specific periods: from 1 July to 30 September in the Niayes basin and from 1 May to 31 October in the Casamance basin. The remaining months were considered the dry season. Results for the Niayes basin showed significant effects of attractant type, season, and their interaction (all $p < 0.001$). Model predictions indicated that the largest difference between attractants occurred during the wet season, when the Econex attractant yielded substantially lower catches than the Biosystemes attractant (approximate back-transformed fitted mean catches of 178 and 479, respectively). During the dry season, although the difference was also significant, it was very slight and in favor of the Econex attractant (5.5 vs. 5.3). Results were largely similar in the Casamance basin with Econex being less efficient than Biosystemes during the the wet season (approximate back-transformed fitted mean catches of 225 and 414, respectively). During the dry season, efficiency was slightly higher for the Biosystemes than the Econex attractant (87 vs. 78, respectively). Due to the marked difference in attractant efficiency during the wet season, we used the maximum catch among the three traps deployed in each orchard, rather than the mean catch, to construct the abundance time series.

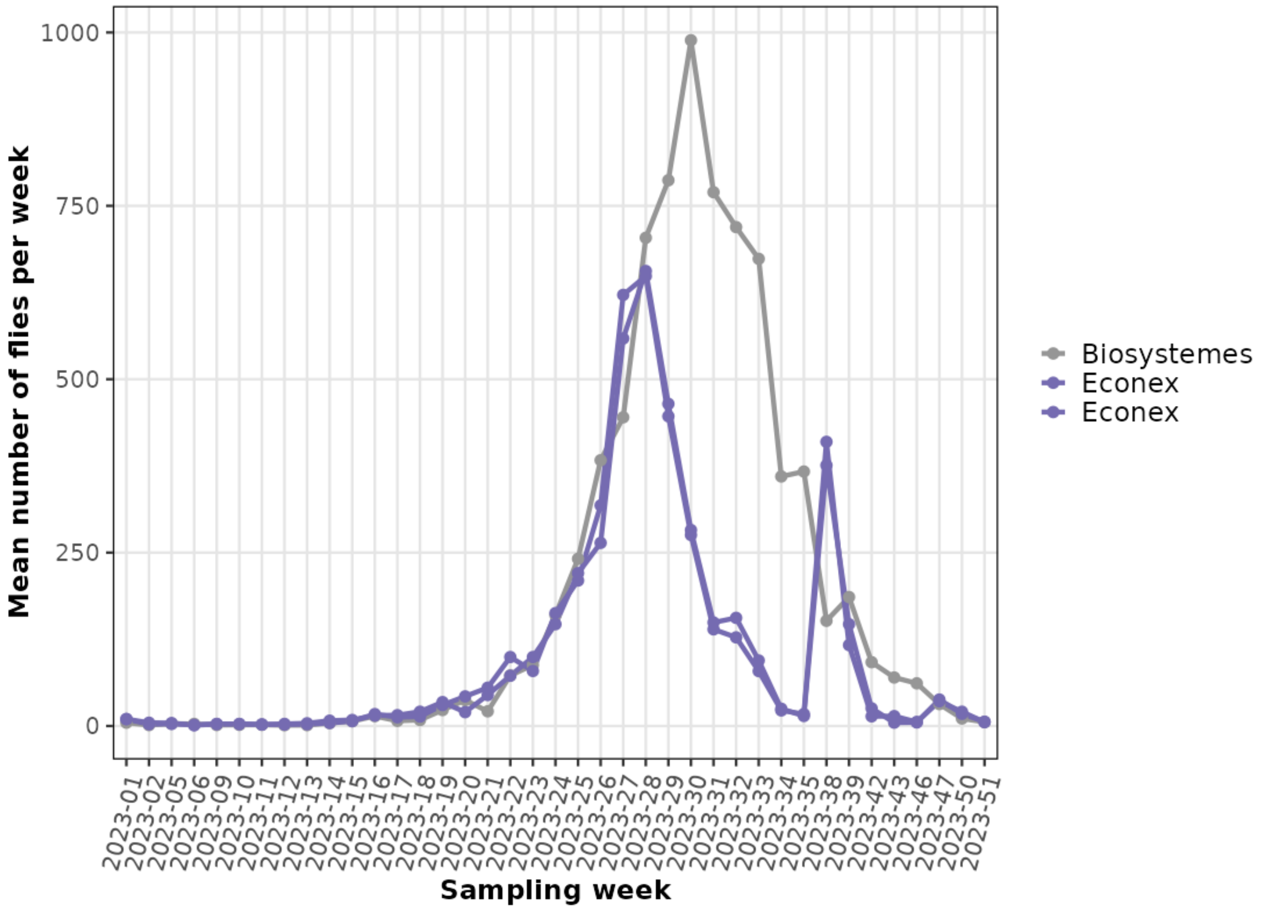


**Figure S1: Methyl-eugenol attractant efficiency.** The mean number of flies trapped per week in 2023 over the 28 orchards in the Niayes is displayed according to the brand of the attractant used in the 3 traps set up within each orchard)

### Supplementary Material S2 – Bayesian estimation of the *eps* and $t_0$ parameters from *B. dorsalis* abundance time series, using a modified version of the POPFIT model

In this work, we used the mechanistic model POPFIT developed by Soulsby and Thomas (2012) to model population curves and estimate demographic parameters (the start date of population growth $t_0$ and the mean level of abundance outside of the mango season *eps*), from *B. dorsalis* annual abundance time series collected in two Senegalese mango production basins, the Niayes and Casamance.

#### 1 - Modification of the POPFIT model to account for non-zero abundance outside the demographic peak

The initial POPFIT model (1), firstly applied to butterfly populations, allows to estimate four biological parameters: the start date of the eclosion period, which can be considered as the start date of the annual population growth ($t_0$), the total population ($N$), the lenght of the eclosion period ($T_E$) and the mean life span ($T$). In Senegal, *B. dorsalis* populations exhibit annual demographic kinetics consistent with the proposed model framework: a rapid increase in catches at the beginning of the mango season until reaching a peak in abundance, followed by a strong decrease in population size, similarly to the dynamics of butterfly populations studied in Soulsby and Thomas (2012). As shown in (2), the initial POPFIT model assumes a null population size when $t < t_0$ (i.e., before the onset of population growth), which aligns well with *B. dorsalis* dynamics in the Niayes basin (Caumette et al. 2024). However, this assumption does not reflect the kinetics observed in the Casamance basin, where flies are present year-round. Therefore, we modified the POPFIT model to relax the zero-abundance assumption before $t_0$ by adding the parameter *eps*, which estimates the average abundance outside the mango season (3).

The POPFIT model is initially formulated as:

$$\frac{dn}{dt} = \frac{N}{T_E} E(t'/T_E) - \frac{n}{T} \quad (1)$$

with $t' = t - t_0$.

Using a sine-cubed function for the reproduction function $(1/T_E)E(t'/T_E)$ (Soulsby and Thomas 2012), the explicit formula for $n(t)$ is:

$$\begin{cases} n(t) = 0 \text{ for } t < t_0 \\ n(t) = \frac{3N}{4(a^2+9)} \left[ asin^3X - 3sin^2X.cosX + \frac{6}{a^2+1}(asinX - cosX + e^{-aX}) \right] \text{ for } t_0 \leq t \leq t_0 + T_E \\ n(t) = \frac{9Ne^{-aX}(1+e^{a\pi})}{2(a^2+9)(a^2+1)} \text{ for } t > T_E + t_0 \end{cases} \quad (2)$$

with $X = \pi t'/T_E$ and $a = T_E/(\pi T)$.

The explicit formula for $n(t)$ in the modified model is then:

$$\begin{cases} n(t) = eps \text{ for } t < t_0 \\ n(t) = \frac{3N}{4(a^2+9)}\left[asin^3X - 3sin^2X.cosX + \frac{6}{a^2+1}(asinX - cosX + e^{-aX})\right] + eps \text{ for } t_0 \leq t \leq t_0 + T_E \\ n(t) = \frac{9Ne^{-aX}(1+e^{a\pi})}{2(a^2+9)(a^2+1)} + eps \text{ for } t > T_E + t_0 \end{cases} \quad (3)$$

with $X = \pi t'/T_E$ and $a = T_E/(\pi T)$.

#### 2 - Bayesian parameter inference

Parameter inference was performed within a mechanistic–statistical framework (Papaïx et al. 2022), independently for each time series (one per orchard and year). As previously done in Caumette et al. (2024), a probabilistic model was defined, conditioned on the theoretical abundance $n$ (see (3)) assuming that, for one year in one orchard, the number of catches at time $t$ followed a Poisson distribution:

$$n_t^{obs}|n(t) \sim Pois(n(t) + 0.01) \quad (4)$$

The Bayesian inference was done using Nimble (de Valpine et al. 2017), with the prior distributions for the model parameters presented in Table S1. We ran 5 MCMC chains, each with 350,000 iterations and a burn-in period of 200,000. After that, the chains were thinned every 150 iterations to obtain 1000 values per chain that were gathered to form the posterior distributions of each parameter.

**Table S1: Parameter prior distributions used to perform the Bayesian inference.** For $t_0$, the minimum and maximum bounds of the uniform distribution were adjusted for each time series when necessary, to improve the fit to the data and the convergence. sd: standard deviation.

| model parameter | distribution | distribution parameters |
|---|---|---|
| $N$ | normal | mean = 8.5, sd = 0.3 |
| $T$ | uniform | min = 1, max = 12 |
| $T_E$ | uniform | min = 1, max = 30 |
| $t_0$ | uniform | between 1 and 35 (adjusted for each time series) |
| $\sigma_{eps}$ | uniform | min = 0, max = 10 |
| $\mu_{eps}$ | normal | mean = 0, sd = 10 |
| $eps$ | log-normal | mean = $\mu_{eps}$, sd = $\sigma_{eps}$ |

Values of the Gelman-Rubin statistic were $< 1.1$ for all estimates of $eps$ and $t_0$ and the visual control of the trace plots of the MCMC chains also showed good convergence.

### Supplementary Material S3 – Construction and preprocessing of environmental predictors

#### 1 - Description of the crop system within orchards

To characterize the cropping system of the 56 monitored orchards, we followed a methodology similar to Caumette et al. (2024), with the additional implementation of a phenological classification of orchard host plants. We characterized each orchard using three main components: (i) the ratio of illuminance between the interior and exterior of the orchard canopy to quantify local canopy density and light penetration (hereafter "lux ratio"); (ii) the presence or absence of agricultural practices potentially affecting early *B. dorsalis* population growth within orchards, including irrigation, soil management, and intercropping with vegetable crops, based on interviews with mango producers; and (iii) the abundance and phenological status of host species and mango cultivars within orchards, as detailed in the following section.

##### 1.1 - Phenological classification

Based on our phenological surveys as well as the literature (see Caumette et al. (2024) and Ndiaye (2009)) and expert knowledge, we grouped the various host trees and mango cultivars into phenological categories depending on the period of fruit availability, while accounting for their frequency. Thus, the most frequent mango cultivars in each basin were considered as different categories while less common cultivars were divided in two groups based on their phenology, early cultivars (M1) and late cultivars (M2). With the same reasoning, potential alternative host species were grouped into three phenological categories: *AH1* includes citrus species that produce fruit from December to April, outside of the mango season; *AH2* includes species with fruit availability all year round; *AH3* includes species that produce fruit mostly during the mango season, i.e. from April to November. Then, for each monitored orchard, we calculated the proportion of each category based on survey data (see Table S2 and Table S3 for averaged proportions across orchards in the Niayes and Casamance basins, respectively).

**Table S2: Distribution of 79,314 trees across eight phenological classes in the 28 monitored orchards in the Niayes.** Main mango cultivars were considered as distinct phenological classes, whereas other mango cultivars (M) and alternative host plants (AH) were classified separately into early or late phenological classes (1–3). The number and proportion of trees are reported for each class.

| Type | Phenological class | Taxonomic name | Period of fruit availability | Total number | Proportion |
|---|---|---|---|---|---|
| mango | Mkent | *Mangifera indica* cv. Kent | June-July to August | 65126 | 82 % |
| | Mbdh | *Mangifera indica* cv. Boukodiekhal (BDH) | April to September | 1251 | 1.6 % |
| | Mkeitt | *Mangifera indica* cv. Keitt | July to October | 7304 | 9.2 % |
| | M1 (early cultivars) | *Mangifera indica* cv. Sewe, Papaye, Greffal, Dieg bou gatt (Dbg), Birane Diop | April to May-June | 247 | 0.5 % |
| | M2 (late cultivars) | *Mangifera indica* cv. Solom, Simiki, Ronde, Palmer, Bambe, Amelie | June to August | 43 | 0.1 % |
| alternative hosts | AH1 | *Citrus X limon, Citrus reticulata, Citrus maxima* | December to April | 4603 | 5.8 % |
| | AH2 | *Carica papaya, Persea americana, Musa spp, Manilkara zapota* | potentially all year round | 351 | 0.4 % |
| | AH3 | *Cola spp, Anacardium occidentale, Psidium guajava, Annona spp* | April to November (during or after the mango season) | 389 | 0.5 % |

**Table S3: Distribution of 7,537 trees across eight phenological classes in the 28 monitored orchards in Casamance.** Main mango cultivars were considered as distinct phenological classes, whereas other mango cultivars (M) and alternative host plants (AH) were classified separately into early or late phenological classes (1–3). The number and proportion of trees are reported for each class.

| Type | Phenological class | Taxonomic name | Period of fruit availability | Total number | Proportion |
|---|---|---|---|---|---|
| mango | Mkent | *Mangifera indica* cv. Kent | May-June to September | 1518 | 20.1 % |
| | Mkeitt | *Mangifera indica* cv. Keitt | May-June to September | 2617 | 34.7 % |
| | Mdiourou | *Mangifera indica* cv. Diourou | May-June to July | 421 | 5.6 % |
| | M1 (early cultivars) | *Mangifera indica* cv. Sucar, Pomme, Papaye, Dieg bou gatt (Dbg), Boukodiekhal (BDH), Birane diop | April to July | 247 | 3.3 % |
| | M2 (late cultivars) | *Mangifera indica* cv. Sierra Leone, Peche, Longue, Amoulene | May-June to August | 336 | 4.5 % |
| alternative hosts | AH1 | *Citrus sinensis, Citrus X limon, Citrus reticulata* | December to April | 2106 | 27.9 % |
| | AH2 | *Carica papaya* | potentially all year round | 280 | 3.7 % |
| | AH3 | *Psidium guajava* | April to November (during or after the mango season) | 12 | 0.2 % |

#### 2 - Characterization of the surrounding landscape

To characterize the effect of the landscape features surrounding the monitored orchards, we followed the methodology used in Caumette et al. (2024) with two main improvements.

##### 2.1 - Development of a fruit fly dedicated land cover map

First, we developed a dedicated typology of 16 land cover classes (Table S4), using a classification approach based on a convolutional neural network applied on time series (January 2022 to September 2023) of satellite imagery Sentinel-1, Sentinel-2 along with an elevation layer (SRTM). For each pixel, the model provides a confidence score from 0 to 1 for each possible landscape class. This score informs on the uncertainty in the classification of the pixels, which may be important for landscape features that are difficult to identify. The production framework and the R and Python scripts used to generate this landscape typology are currently under review as part of a data paper and will be made publicly available through the data paper if it is accepted for publication.

**Table S4: Description of the 16 classes of the land cover typology**

| Land use class | Description |
|---|---|
| Lowland irrigated crop | agricultural areas equipped with permanent irrigation infrastructure (e.g., pumps, canals, boreholes), enabling cultivation during both wet and dry seasons |
| Irrigated farming | Irrigated agricultural plots, excluding lowlands, which can be cultivated during both the wet and dry seasons. |
| Rainfed crop | agricultural plots, mainly vegetable, located in the lowlands and watered using on-site water resources, typically without formal irrigation infrastructure |
| Water | water area, including rivers, lakes, sea water |
| Dense forest | wooded area with high tree cover (typically, the tropical forest of Basse-Casamance) |
| Dry forest | wooded area with low to moderate tree cover |
| Mangrove | evergreen, salt-tolerant woody vegetation, including trees and shrubs, rooted in brackish or saline water |
| Wetland | land saturated with water, either permanently or seasonally, whether cultivated or not (e.g. rice field, swamp) |
| Shrub savannah | mix of scattered shrubs and grasses |
| Herbaceous savannah | land dominated by grasses and other non-woody plants, with few or no shrubs or trees |
| Floodable bare ground | bare ground subject to seasonal flooding during the rainy season or influenced by tidal fluctuations |
| Bare ground | area of bare soil, including sand, rock, quarry |
| Dense urban | highly urbanized area (e.g. city) |
| Scattered urban | area where buildings or settlements are dispersed (typically found in rural or peri-urban areas) |
| Orchard | mango (in majority) and Citrus spp. plantations |
| Cashew orchard | cultivation of cashew trees, mainly found on the border between Senegal and Guinea-Bissau |

##### 2.2 - Estimation of the spatial influence of land cover classes with *scalescape*

Second, we used the R package *scalescape* v. 0.0.0.9 (Lowe et al. 2022) to estimate the spatial range of influence of the landscape features on the demographic parameters $t_0$ and *eps*. This package implements a similar framework to *siland* (Carpentier and Martin 2021), used in Caumette et al. (2024), but additionally allows the use of quantitative landscape values (here, confidence scores assigned to each pixel of the landscape map) and enables the joint estimation of spatial influence functions (SIFs) for all landscape classes within a single model. For each basin and each parameter ($t_0$ and *eps*), the sampling year was included as a local explanatory variable in the *scalescape* model, which was run independently on 50 out of the 500 sample sets to limit computation time and energy consumption. We considered only the sample sets with a significant effect (p-value $< 0.01$) to compute the median of the *SIF* ranges over all landscape classes. Then, for each parameter and basin, we considered the median value of the *SIF* and a maximum distance of 6000 m, to derive the final Gaussian weighting function used to describe the decay of landscape influence with distance from each sampling site (see Lowe et al. 2022). The results are shown in Figure S2. The median values of the SIF were: 600 m and 500 m for the *eps* parameter and 660 m and 790 m for the $t_0$ parameter in the Niayes and Casamance basins, respectively.

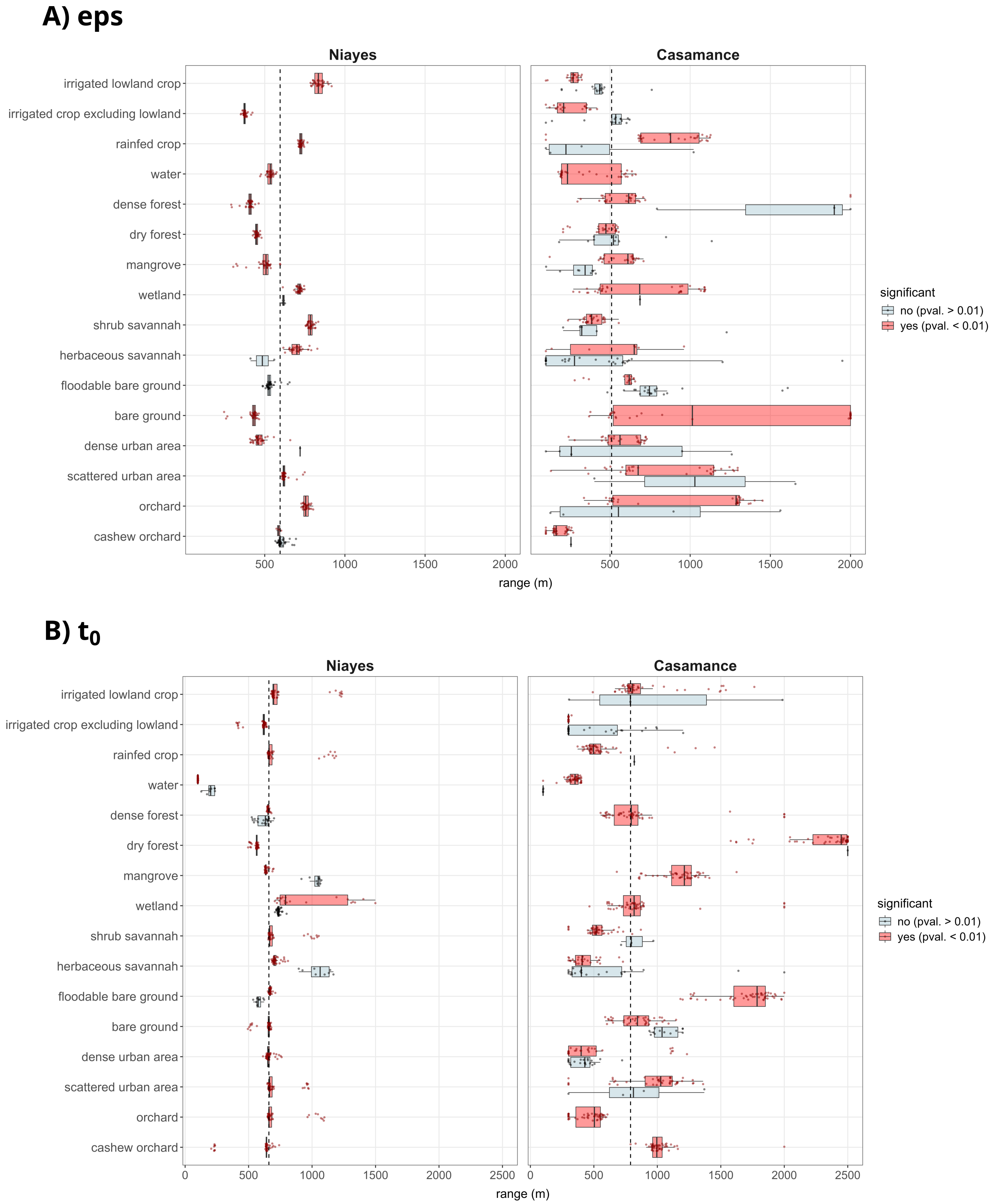


**Figure S2: Scalescape results.** For each demographic parameter *eps* (A) and $t_0$ (B), the figure shows the distribution of the range of the spatial influence function estimated independently on 50 samples sets. Vertical dashed lines represent the median values over all landscape classes.

#### 3 - Data sources for weather and vegetation

The spatio-temporal variability of weather and vegetation features was analysed over the production basins and the period of sampling using different data sources. First, monthly precipitation estimates were retrieved from the Tropical Applications of Meteorology using SATellite data and ground-based observations (TAMSAT) at 4 km resolution (Maidment et al. 2014; Maidment et al. 2017; Tarnavsky et al. 2014). Second, MODIS data were used to retrieve day-time land surface temperatures (MOD11A2) at 1 km resolution and vegetation indices (MOD13Q1- 16-Day L3 Global 250m; Didan 2015; Didan et al. 2015): the Enhanced Vegetation Index (EVI), which reflects changes in canopy density and structure, and the Normalized Difference Water Index (NDWI), an indicator for vegetation water content (Gao 1996; Gu et al. 2007) that we calculated according to the formula of NDWI 2130 defined in Chen et al. (2005). Raster of physical estimates (temperature, precipitation, EVI, NDWI) were aligned to a final 1 km spatial resolution. Then, we considered the rasters for the months of January, February and March, which correspond to the main period of low abundance of *B. dorsalis*, and the different estimates were averaged (day-time temperatures and EVI and NDWI vegetation indices) or summed (precipitation) over these three months.

### Supplementary Material S4 – Model testing in the GP-Boost framework

#### 1 - Random effect models

GPBoost allows random effects to be modelled as grouped effects (potentially nested) and/or Gaussian processes (Sigrist 2022). Here, we tested several random effects models, using the function `GPModel` of the R package *gpboost*. All models were tested independently on each sample set for the parameters *eps* and $t_0$, with the later being log-transformed.

**Model 1: Grouped random effects**

```
mod1 <- GPModel(group_data=year, likelihood="gaussian")
```

**Model 2: Spatial Gaussian process random effects with every group having the same spatial effect**

```
mod2 <- GPModel(gp_coords=xycoords, likelihood="gaussian")
```

**Model 3: Spatial Gaussian process random effects with every period having a different spatial effect**

```
mod3 <- GPModel(gp_coords=xycoords, cluster_ids=as.vector(year), likelihood="gaussian")
```

**Model 4: Grouped period and site random effects and a spatial Gaussian process for residual spatial correlation**

```
mod4 <- GPModel(group_data=year, gp_coords=xycoords, likelihood="gaussian")
```

**Model 5: approximated spatio-temporal Gaussian process model**

```
mod5 <- GPModel(gp_coords=as.matrix(cbind(xycoords,as.numeric(year))), likelihood="gaussian")
```

#### 2 - Hyperparameter tuning

Hyperparameters of the GPBoost algorithm were optimised using a grid search procedure based on a 4-fold cross validation (function "gpb.grid.search.tune.parameters" of the *gpboost* R package) performed independently on each sample set. The following hyperparameter values were considered:

- learning rate (learning_rate): 0.01, 0.05, 0.1, 0.5, 0.8, 1
- minimum data in leaf (min_data_in_leaf): 2, 3, 5, 10, 20
- maximum depth of trees (max_depth): 2, 3, 5, 8, 10, 20, 30, 40

The number of iterations (i.e. the number of trees), best_iter, was automatically optimised during the tuning step, with a maximum number of iterations possible set to 2000 and the early_stopping_rounds parameter to 5, i.e. the process stops if the model's performance on the validation set does not improve for 5 consecutive iterations. For each parameter combination, the Mean Square Error (MSE) was calculated. Then, for each tested model, the lowest MSE and the associated hyperparameter values were retained.

#### 3 - Selection and training of the best model

Based on the cross-validation results (MSE), the best model corresponded to the model including a grouped year random effect. This model was then trained independently on each sample set using the combination of hyperparameter values that minimised the MSE for the corresponding sample set.

### Supplementary Material S5 – GPBoost predictive performance

Performance of the GPBoost models were assessed by predicting *eps* and $t_0$ in each basin, considering either all predictors or the top-ranked predictors. Each sample set was randomly split into a training and a test dataset, i.e. 80% and 20% of the data, respectively. Model accuracy was then assessed by computing the Pearson correlation coefficient between predicted and observed values as well as the Root Mean Square Error (RMSE) of the models for each sample set. The results are shown Figure S3 (*eps* parameter) and Figure S4 ($t_0$ parameter).

**A)**

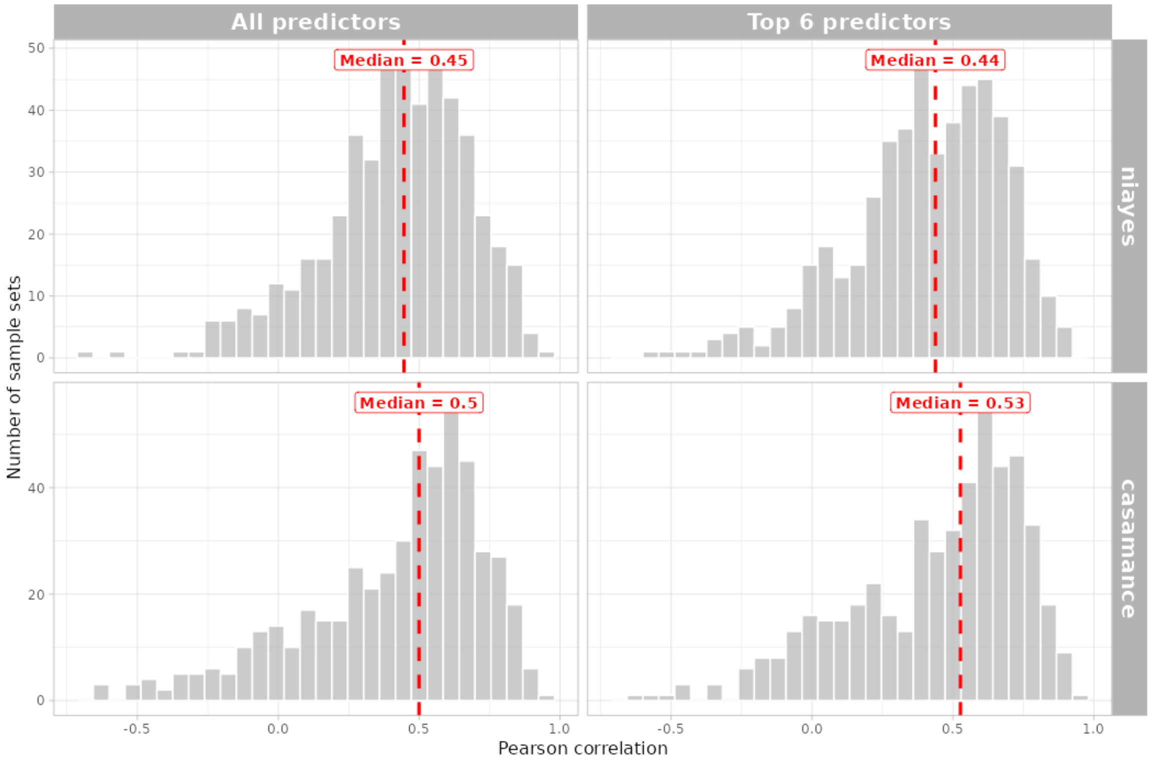


**B)**

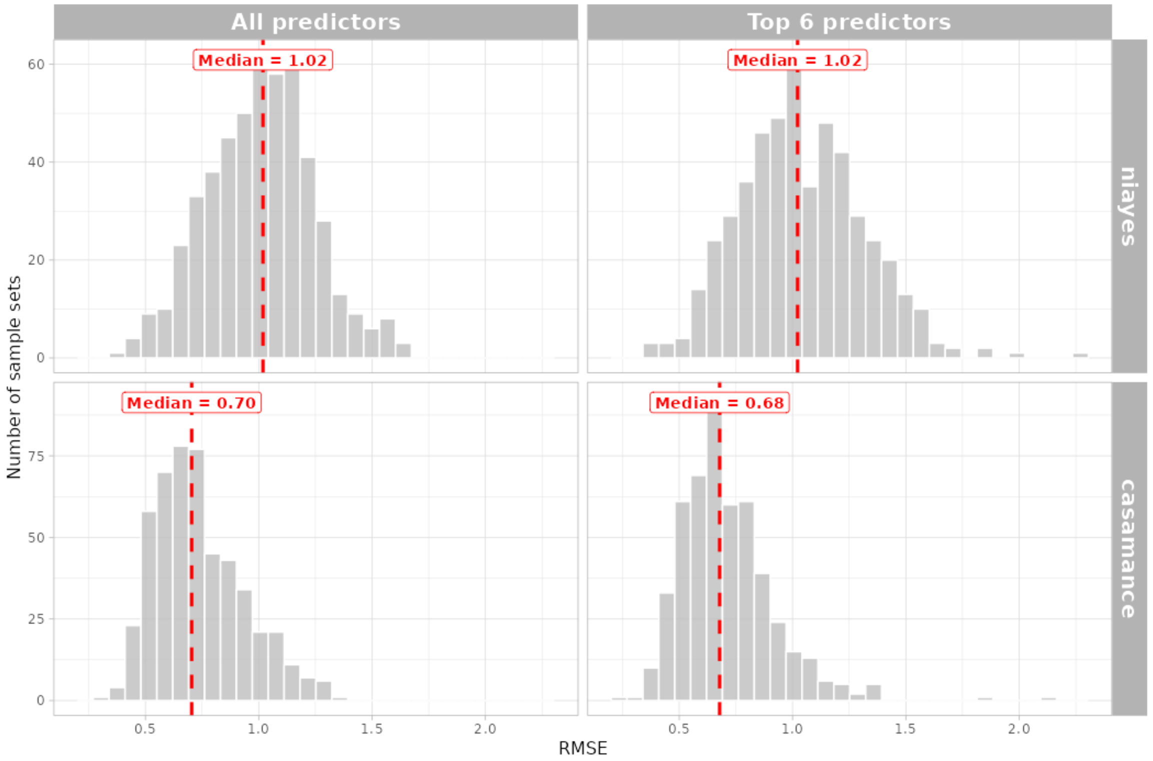


**Figure S3: Predictive performance of the GPBoost model on the *eps* parameter.** For each sample set, the training and prediction with GPBoost were made respectively on 80% and 20% of the data, using the model including all predictors (left panel) or the model including only six top-ranked predictors (right panel), separately in each basin (Niayes and Casamance). The quality of the prediction was evaluated by calculating (A) the Pearson correlation coefficient and (B) the Root Mean Square Error (RMSE) for the 500 sample sets. For each model, the vertical line and the associated number correspond to the median value of the variable (RMSE or Pearson coefficient) over the 500 sample sets.

**A)**

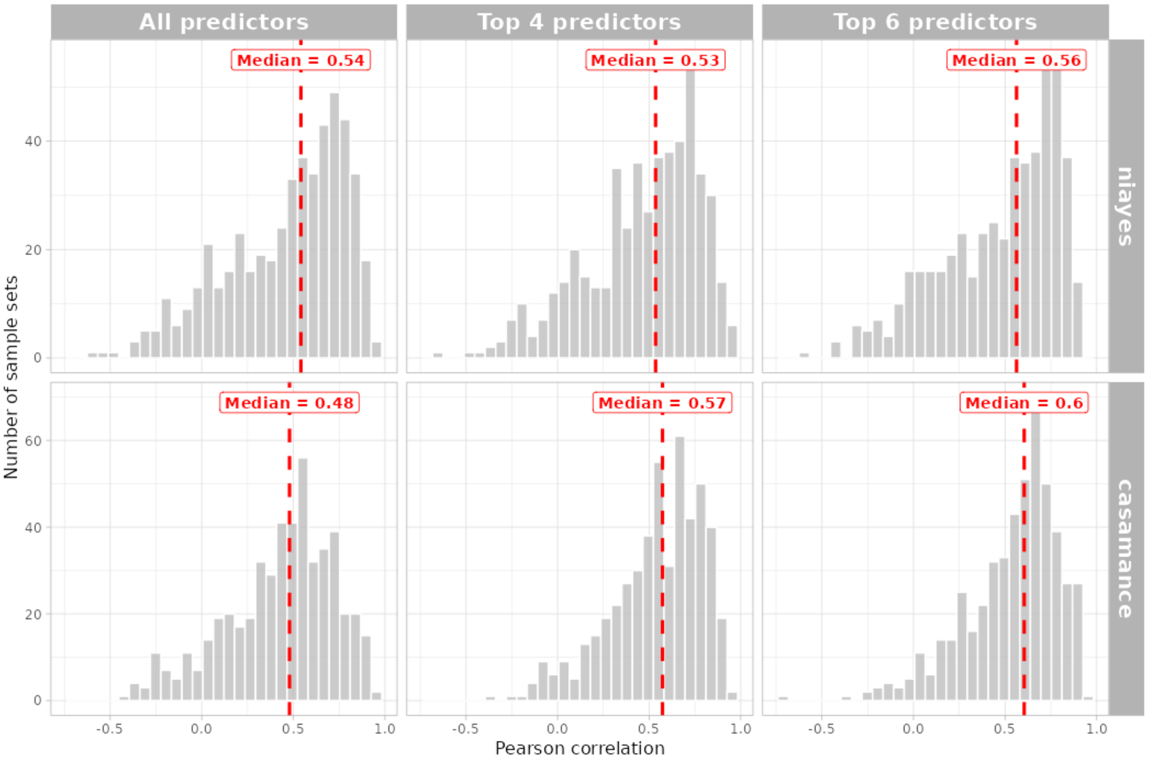


**B)**

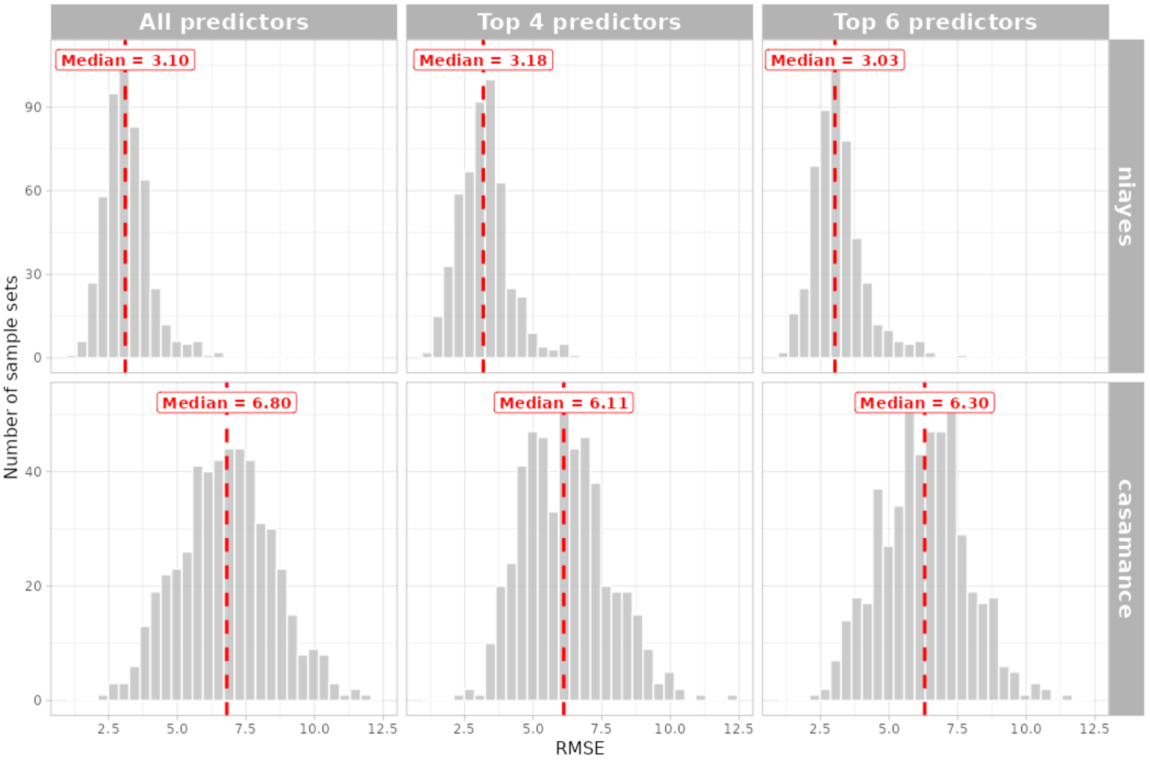


**Figure S4: Predictive performance of the GPBoost model on the $t_0$ parameter.** For each sample set, the training and prediction with GPBoost were made respectively on 80% and 20% of the data, using the model including all predictors (left panel) or the models including either four or six top-ranked predictors (middle and right panels), separately in each basin (Niayes and Casamance). The quality of the prediction was evaluated by calculating (A) the Pearson correlation coefficient and (B) the Root Mean Square Error (RMSE) for the 500 sample sets. For each model, the vertical line and the associated number correspond to the median value of the variable (RMSE or Pearson coefficient) over the 500 sample sets.
